# MTG2: An efficient algorithm for multivariate linear mixed model analysis based on genomic information

**DOI:** 10.1101/027201

**Authors:** S.H. Lee, J.H.J. van der Werf

**Affiliations:** School of environmental and Rural Science, University of New England, NSW 2351, Australia; Queensland Brain Institute, The University of Queensland, QLD 4072, Australia.

## Abstract

We have developed an algorithm for genetic analysis of complex traits using genome-wide SNPs in a linear mixed model framework. Compared to current standard REML software based on the mixed model equation, our method could be more than 1000 times faster. The advantage is largest when there is only a single genetic covariance structure. The method is particularly useful for multivariate analysis, including multitrait models and random regression models for studying reaction norms. We applied our proposed method to publicly available mice and human data and discuss advantages and limitations.

**Availability:** MTG2 is available in https://sites.google.com/site/honglee0707/mtg2.

**Supplementary information:** Supplementary data are available.

## 1. Introduction

Previously, methods have been developed to estimate genetic variance and genetic correlations between complex traits explained by genome-wide SNPs using linear mixed models (Lee, et al., 2012; Maier, et al., 2015; Yang, et al., 2011). Because genetic relatedness among (conventionally) unrelated subjects could be estimated based on genomic information rather than familial relatedness, the model allows estimating genetic effects to be much less confounded with family environmental effects. For the same reason, this approach has also been proposed as a more powerful tool to detect genotype-environment interaction (G × E) (Lee, et al., 2015). That is, in presence of G × E, the genetic correlation between genetic effects in different environments is significantly lower than one (Falconer and Mackay, 1996). In order to capture G × E across a trajectory of multiple environments, random regression models have been proposed for evolutionary and livestock genetics (Kirkpatrick, et al., 1990; Meyer and Hill, 1997). The random regression model is also known as reaction norm model (Kirkpatrick and Heckman, 1989).

In estimating genetic variance explained by genetic markers, Lee and van der Werf (2006) introduced an efficient average information (AI) algorithm to obtain residual maximum likelihood (REML) estimates. As opposed to using Henderson’s mixed model equation (MME) the algorithm was based on using the variance covariance matrix of phenotypic observations directly, hence the term ‘direct AI algorithm’. The algorithm is particularly advantageous when using a dense covariance matrix, such as the genomic relationship matrix (GRM), and with a large number of multiple random effects. The direct AI algorithm is implemented in GCTA-GREML (Lee, et al., 2012; Yang, et al., 2013; Yang, et al., 2011) and MultiBLUP (Speed and Balding, 2014) that has been widely used in human, evolutionary and livestock genetics.

Here, we combine the direct AI algorithm with an eigen-decomposi-tion of GRM, as first proposed by Thompson and Shaw (1990). We apply the procedure to analysis of real data with univariate, multivariate and random regression linear mixed models with a single genetic covariance structure, and demonstrate that the computation efficiency can increase by > 1,000 fold compared with standard REML software based on MME.

## 2. Methods

### 2.1. Model

We used multivariate linear mixed models and random regression models to estimate genetic variances and covariances across multiple traits and among traits expressed in different environments. A linear mixed model can be written as

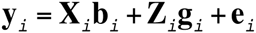

where **y***_i_* is a vector of trait phenotypes, **b***_i_* is a vector of fixed effects, **g***_i_* is a vector of additive genetic value for individuals and **e***_i_* are residuals for trait or environment *i*. **X** and **Z** are incidence matrices. More details can be found in the Supplementary Note. To model genotype-environment interactions, a random regression model attempts to fit effects as a function of continuous variable (Kirkpatrick, et al., 1990; Meyer and Hill, 1997) as

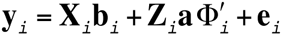

where **a** is a n (# records) by *k* matrix of genetic random regression coefficients, **Φ**_i_ is the *i*th row in a *p* by *k* matrix of Legendre polynomials evaluated for *p* points on the trajectory, and *k* is the order of Legendre polynomials. This model is explicitly described in the Supplementary Note. The genetic covariance structure was constructed based on genome-wide SNPs.

### 2.2. Algorithm

REML is often solved using Newton-Raphson or Fisher’s scoring method where variance components are updated based on observed (Hessian matrix) or expected second derivatives of the log likelihood (Fisher information matrix). In order to increase the computational efficiency in obtaining REML estimates, Gilmour et al. (1995) used the average of the Hessian and Fisher information matrix that was estimated based on Henderson’s mixed model equation (MME). The MME-based average information (AI) algorithm is efficient particularly when the genetic covariance structure fitted in the model is sparse. When using dense covariance structures such as GRM, the computational efficiency of a direct AI algorithm is substantially more efficient that the MME-based AI algorithm (Lee and Van der Werf, 2006). Here, we extend the direct AI algorithm by implementing an eigendecomposition of the genetic covariance structure as proposed by Thompson and Shaw (1990).

In recent studies the eigen-decomposition technique has been implemented in a Newton-Raphson algorithm in univariate and multivariate linear mixed models (Zhou and Stephens, 2014). We show that an implementation in the direct AI algorithm is mathematically straightforward and is computational more efficient especially in multivariate linear mixed models (Supplementary Note). Moreover, we show how our proposed algorithm can be efficiently applied to a random regression model (Supplementary Note).

### 2.3. Data

We used heterogeneous stock mice data (http://mus.well.ox.ac.uk/mo-use/HS/) to estimate genetic variances and covariances of complex traits explained by genomewide SNPs. After a stringent QC of genotypic data, we used 9,258 autosomal SNP from 1,908 individuals. We used phenotypes of four glucose values (taken at 0, 15, 30 and 75 minutes after intraperitoneal glucose injection in a model of type 2 diabetes mellitus) as well as body mass index (BMI). We analysed this data in a five-trait linear mixed model. We also applied a random regression model for the repeated glucose measures.

Secondly, we used human data from the Atheroscierosis Risk in Communities (ARIC) cohort (psh000280.v3.p1) (Sharrett, 1992). A similar stringent QC as above was applied to the available genotypes. In addition, we randomly removed one of each highly related pair of relatedness > 0.05 to avoid bias due to population structure or family effects. After QC, 7,263 individuals and 583,058 SNPs remained. We used BMI, triceps skinfold (TS), waist girth (WG), hip girth (HG), waist to hip ratio (WHR), systolic blood pressure (SP), diastolic blood pressure (DP) and hypertension (HP) that were fitted in an eight-trait linear mixed model.

Missing phenotypic values were less than 10% and 1% for each trait for the mice and the human data, respectively. They were imputed with their expected values from univariate linear mixed model fitting each trait separately.

### 2.4. Software

We implemented the direct AI algorithm and the eigen-decomposition technique into software named as MTG2. We compared MTG2 with GEMMA (Zhou and Stephens, 2014), ASReml (Gilmour, et al., 2006) and WOMBAT (Meyer, 2007). GEMMA uses the eigen-decomposition technique in the Newton-Raphson algorithm. ASReml and WOMBAT are well known REML software that use an MME-based AI algorithm.

## 3. Results

When using WTCCC heterogeneous mice data (N=1,908) for multivariate linear mixed model with up to five traits, MTG2 only took a few seconds, which was a few thousands times faster than ASReml and WOMBAT and few times faster than GEMMA (Table 1). Estimated SNP-heritability and genetic correlations between traits are shown in Table S1. REML parameters after convergence were essentially the same between different software, as shown in Table S8 and S9.

**Table 1.** Computing time for each software running at 2.7 GHz CPU when using WTCCC heterogeneous stock mice data (N=1908).

|  | MTG2 | GEMMA | ASReml | WOMBAT |
| --- | --- | --- | --- | --- |
| # traits | Multivariate linear mixed model |  |  |  |
| 1 | 1 sec | 1 sec | 2 min | 17 sec |
| 3 | 1 sec | 1 sec | 210 min | 9 min |
| 5 | 2 sec | 6 sec | 950 min | 60 min |
| # order | Random regression model |  |  |  |
| 1 | 2 sec | NA<sup>a</sup> | 4 min | 3 min |
| 2 | 2 sec | NA | 82 min | 30 min |
| 3 | 2 sec | NA | 310 min | 54 min |

<sup>a</sup>GEMMA does not have a function for the random regression model.
For MTG2 and GEMMA, it took ~4 seconds for the eigen-decomposition, which only has to be done once per dataset then can then be reused for multiple analyses.

When using a random regression model, the computing time for MTG2 was a few seconds, which did not change with increasing the order of the fit in the model (Table 1). However, the computational efficiency of ASReml or WOMBAT was lower and computing time increased substantially with increasing the order of the fit (Table 1). GEMMA does not have a function for random regression models. The estimated results from the random regression model are described and shown in Supplementary data (Table S2 and Figure S1).

When using the ARIC cohort human data (psh000280.v3.p1), the pattern of the computing time was similar to that for the heterogeneous mice in that MTG2 and GEMMA performed similar although MTG2 became relatively faster with increasing number of traits (Table S4). ASReml and WOMBAT were too slow to run for this data set. Table S6 shows estimated SNP-heritability and genetic correlations between obesity and blood pressure traits.

## 4. Discussion

There are two limitations to MTG2 as well as GEMMA. The eigen-decomposition technique cannot be used with more than one GRM as also noted in Zhou and Stephens (2014) unless a special condition is satisfied, i.e. one full-rank GRM and multiple low-rank GRMs are given (Speed and Balding, 2014). In models with multiple GRMs, GEMMA cannot be used and MTG2 becomes slow although it is still considerably faster than ASReml and WOMBAT (Table S5). Secondly, the eigen-decomposition technique requires a balanced design (i.e. no missing phenotypes across traits). Phenotypic imputation can be used for phenotypic missing values. We used imputed missing phenotypes for the mice data (<10% missing for each trait) although MTG2 without the eigen-decompostion could still be used for the data including missing values. We observed that the results from the data with and without imputing missing phenotypes were not much different (Table S2 and figure S2). For the human data, missing phenotypes were less than 1%, therefore the results with and without imputing missing phenotypes were almost identical (result not shown). Finally MTG2 and WOMBAT can facilitate a parallel computation that increases the efficiency further.

## 5. Implication

There are three novel aspects in this application note. The first and most important is to estimate parameters for the random regression models with the direct AI algorithm. The second is the use of the decomposition in the AI algorithm in a multivariate model. The third is to use of the decomposition in the AI algorithm in the random regression models. MTG2 can be used for a wider range of statistical models than GEMMA, including multivariate linear mixed models, random regression models and multiple random effects models. GEMMA can only be used for a single random effect model in multivariate linear mixed models (Table S7). For random regression models or/and multiple random effects models, the computational efficiency for MTG2 (even without the eigen-decomposition) is considerably higher than that of ASReml and WOMBAT (Table 1, Table S5 and S7). Therefore, MTG2 can be a useful and efficient tool for complex traits analyses including estimating genetic variance and covariance and G × E.

## Acknowledgements

This study makes use of publicly available data from Wellcome Trust Centre (http://mus.well.ox.ac.uk/mo-use/HS/) and from the database of Genotypes and Phenotypes (dbGaP) under accession psh000280.v3.p1 (see Supplementary Acknowledgements for the full statement).

## Supplementary data

**Availability**: MTG2 is available in https://sites.google.com/site/honglee0707/mtg2.

**Supplementary information**: Supplementary data are available.

### 1. Supplementary Note

#### 1.1. Multivariate linear mixed model

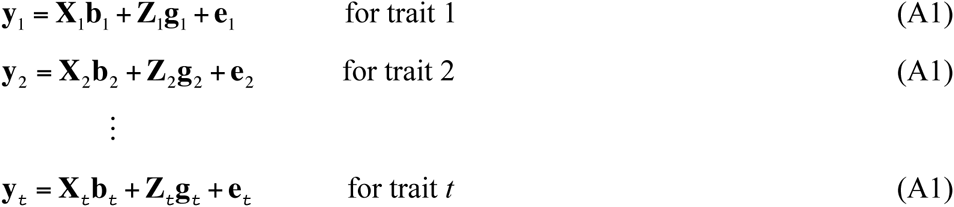

where **y***_i_* is a vector of trait phenotypes, **b***_i_* is a vector of fixed effects, **g***_i_* is a vector of additive genetic value for individuals and **e***_i_* are residuals for trait *i* (*i* = 1, …, *t*). The random effects (**g***_i_* and **e***_i_*) are assumed to be normally distributed with mean zero. **X** and **Z** are incidence matrices. The variance covariance matrix of all observations can be written as,

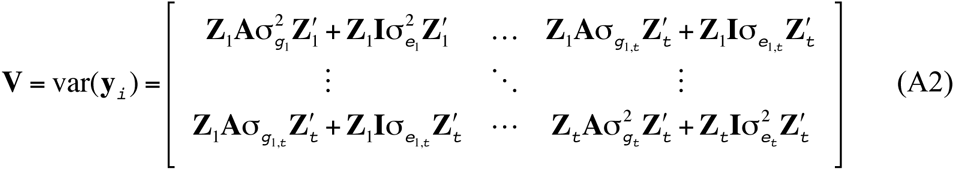

where **A** is the genomic relationship or similarity matrix based on SNP information, and **I** is an identity matrix. The terms, 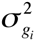 and 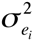 denote the genetic and residual variance of trait *i*, respectively and 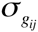 and 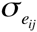 the genetic and residual covariance between traits *i* and *j* (*i*= 1,…, *t, and j*=1,…,*t*).

The log likelihood of the multivariate model is

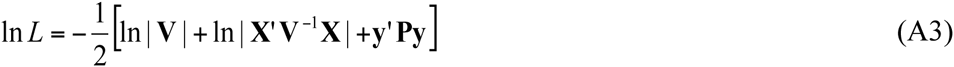

where ln is the natural log, and | | the determinant of the associated matrices. The projection matrix is defined as **P** = **V**^−1^ − **V**^−1^**X**(**X**'**V**^−1^**X**)^−1^**X**'**V**^−1^ with

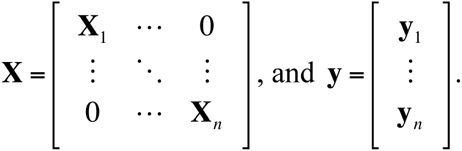

The Newton-Raphson algorithm obtains the residual maximum likelihood (REML) estimates using the following equation (Lynch and Walsh, 1998).

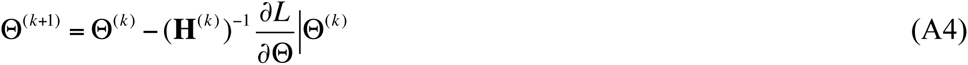

where Θ is a column vector of estimated variance components, *k* is the iteration round, 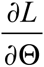 is a column vector of the first derivatives of the log likelihood function with respect to each variance component, and **H** is the Hessian matrix which consists of the second derivatives of the log likelihood function with respect to the variance components. In Fisher’s scoring method, the inverse of the Hessian matrix in (A4) is replaced by its expected value (Lynch and Walsh, 1998) as

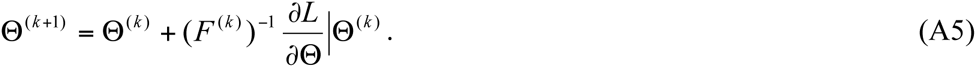

The derivation of the Hessian matrix and the Fisher information matrix has been described in several studies (Lynch and Walsh, 1998; Searle, et al., 1992). The Hessian matrix for the multivariate model is

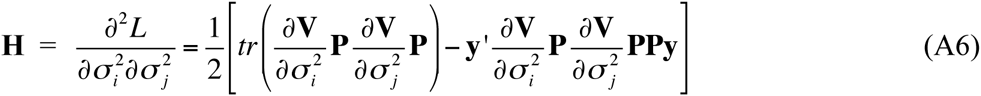

where **y**, **P** and **V** are defined as above. The Fisher information (F) matrix is

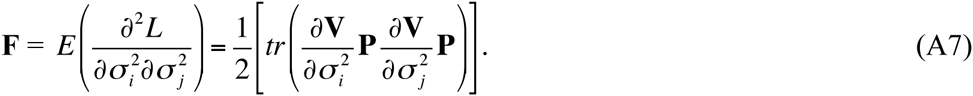

Gilmour et al. (1995) and Johnson and Thompson (1995) used the average of the **H** and **F** that was estimated based on Henderson’s mixed model equation (MME) (Henderson, 1975). Lee and van der Werf (2006) introduced the direct average information algorithm where the average information matrix was derived directly from the **V** matrix. When using a dense genetic covariance structure, this direct average information algorithm is much more efficient than the MME-based average information algorithm. The equation for the iterative AI algorithm is

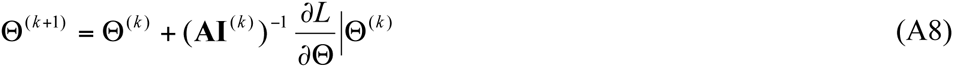

where **AI** is the average information matrix and that for multivariate model can be written as

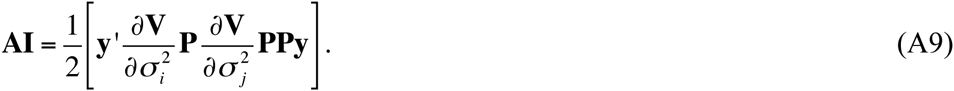

The first derivative for each variance covariance component *i* can be obtained as (Lynch and Walsh, 1998; Searle, et al., 1992)

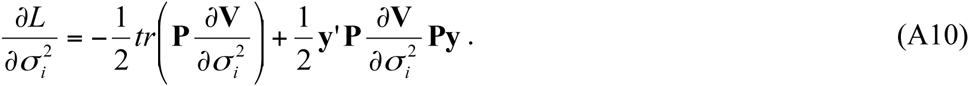

#### 1.2. Increasing computational efficiency

Thompson and Shaw (1990) have proposed an efficient algorithm to obtain the REML estimates using an eigen-decomposition of the genetic covariance matrix. This technique has been used in a number of seminal papers in the context of GWAS (Lippert, et al., 2011; Zhou and Stephens, 2012; Zhou and Stephens, 2014). Lippert et al. (2011) and Zhou and Stephens (2012) have applied it to a univariate linear mixed model using a grid search or the full Newton-Raphson algorithm (i.e. equation A6). Zhou and Stephens (2014) extended their approach to a multivariate linear mixed model framework. Here, we implement the eigen-decomposition technique into the direct AI algorithm (Lee and Van der Werf, 2006) for multivariate REML estimation.

From Thompson and Shaw (1990), the relationship matrix can be decomposed as **A=UDU’** where **I=UU’**. With a linear transform by multiplying the left and right hand side of the multivariate linear mixed model (A1) with **U**, the model can be rewritten as **U**′**y***_i_* **= U**′**X***_i_***b***_i_* **+ U**′**Z***_i_***g***_i_* **+ U**′**e***_i_*. given that the data are a balanced design (i.e. no missing phenotypes). The variance covariance matrix (A2) of the transformed data can be now rewritten as

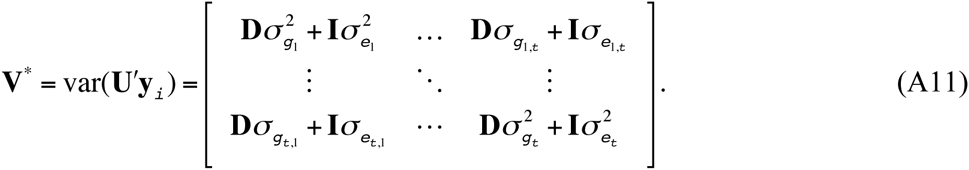

It is also noted that **X** and **y** can be transformed and written as

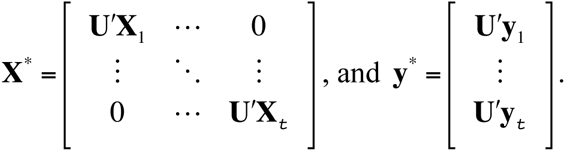

The AI algorithm (A8) needs to evaluate three parts; i) the log likelihood (A3), ii) the **AI** matrix (A9) and iii) the first derivatives (A10). For the evaluation for the log likelihood the inverse **V** is required (A3). Because **V**^*^ (A11) is a matrix with *t*^2^ diagonal blocks, the computational complexity for obtaining the inverse and determinant of **V**^*^ is only *O*(*nt*^3^) where *t* is the number of traits and *n* is the sample size. The product matrix (**X^*^’V^*−1^X^*^**)^−1^ and |**X^*^’V^*−1^X^*^**| require *O* (*nc*^2^*t*+*c*^3^*t*^3^) where *c* is the number of fixed effects for each trait. Following Lynch and Walsh (1998), The product of **P**^*^ and **y**^*^ can be efficiently obtained as

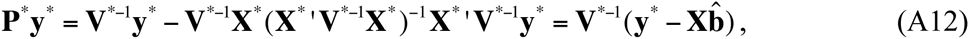

which requires only *O*(*nt*^2^) because also **V**^*−1^ has a simple structure with diagonal blocks. Further, **y^*^P^*^y^*^** requires additional *O*(*nt*).

The second part of the AI algorithm comprises derivation of the **AI** matrix which consists of two terms; 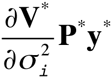 and 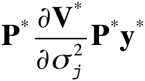. Because the partial derivative, 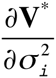 is a partial diagonal matrix with **I**_n×n_ (for **e***_i_*) or **D** (for **g***_i_*) for the *i*th corresponding component in the **AI** matrix, the first term, 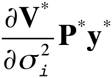 can be easily obtained with *O*(*n*) (note that **P^*^y^*^** is already obtained as above (A12)). Replacing **y^*^** with 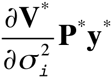 in (A12), the second term can be efficiently obtained as 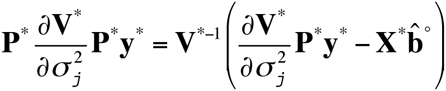 where 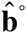 is the generalised least square estimate with 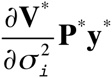 as the response variable (the computational complexity is *O*(*nct*^2^)). Given 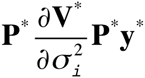 is known, the term, 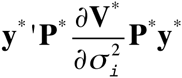 in the first derivative (A10), can be easily obtained with *O*(*n*). The trace term in the first derivative involves only the diagonal part of the **P^*^** and 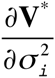, therefore can be efficiently obtained with *O*(*n*). The computational complexity our direct AI algorithm would be ~ *O*(*nt*^3^ + *nct*^6^) within each iteration.

#### 1.3. Random regression linear mixed model

A specific case of a multivariate model arises if the observations with the associated effects can be described as a function of a trajectory, e.g. as a function of time or age. In a random regression model, the model parameters can then be regressed on the trajectory variable, e.g. the genetic covariance structure underlying the data can be modelled as a function of age. If the trajectory variable is an environmental indicator, the random regression model provides a framework to model reaction norms where the genetic effects are modelled as a function of the environmental gradient. This is an elegant form of modelling genotype by environment interaction, or environmental sensitivity of genetic effects. Following Kirkpatrick et al. (1990) and Meyer and Hill (1997), a linear mixed model to incorporate observations at *p* different points in time or in *p* different environments can be written as

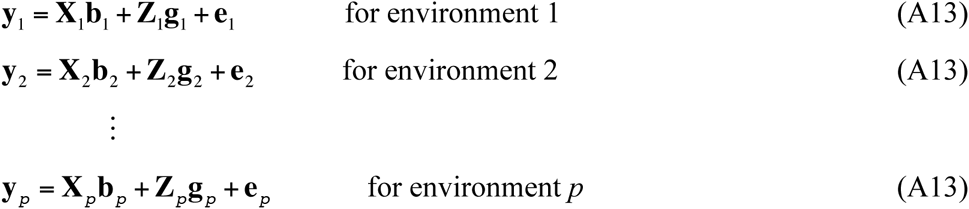

If the environments can be ranked according to a continuous variable on which effects of genes can be regressed, e.g. with an environmental gradient (reaction norm), this can be effectively modelled with random regression coefficients. A random regression model attempts to fit effects as a function of continuous variables, e.g. by using Legendre polynomials (Kirkpatrick, et al., 1990; Meyer and Hill, 1997), as

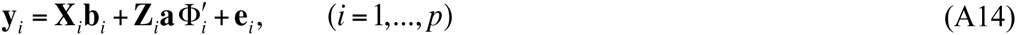

where **a** is a n by *k* matrix of genetic random regression coefficients, **Φ**_i_ is the *i*th row in a *p* by *k* matrix of Legendre polynomials evaluated for *p* points on the trajectory, and *k* is the order of Legendre polynomials. This random regression model reduces a t-dimensional multivariate problem to a k-dimensional problem. The variance and covariance matrix of random regression coefficients is

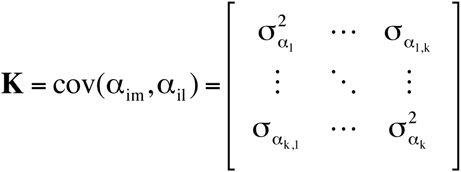

where *α_im_* or *α_il_* is the *m*th or *l*th genetic random regression coefficient for the *i*th individual (*i* = 1 - *n*). The genetic variance and covariance matrix of individual genetic effects across the different environmental conditions is a function of random regression coefficients and Legendre polynomials, and can be written as

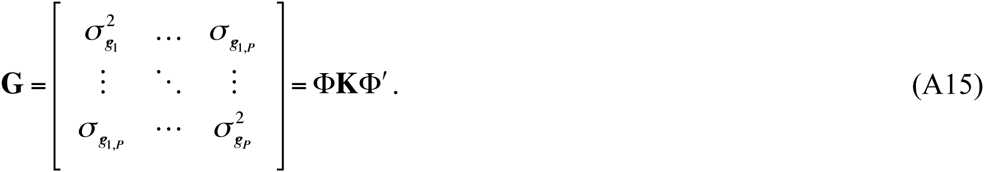

Here we propose the direct average information algorithm for the random regression model, which is suitable for dense genetic covariance structure. The log likelihood of the random regression model is the same as (A3) as

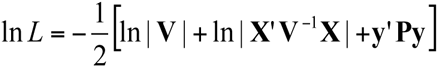

where the matrix **V** can be decomposed as

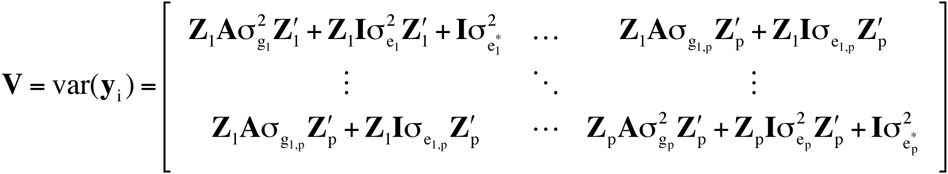

and the matrices **P, X** and **y** are defined as above. From (A15), the additive genetic and permanent environmental effects can be parameterised as a function of Legendre polynomials (**Φ**) and variance covariance matrix of random regression coefficients (**K**) as

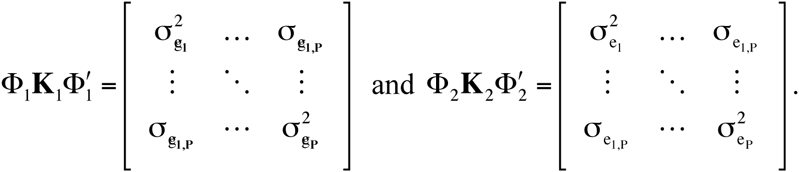

The residual variance within each environment can be stored in a matrix as

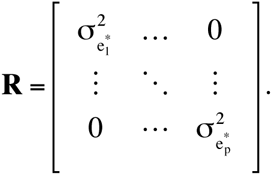

We are interested in obtaining the REML estimate for the elements in the matrices **K**_1_**, K**_2_ and **R**. We use an average information algorithm that requires the first and second derivatives of the likelihood function with respect to each variance component in **K**_1_**, K**_2_ and **R**. The first derivatives and AI matrix can be written as

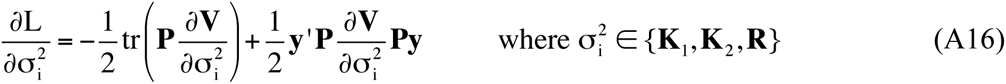

and

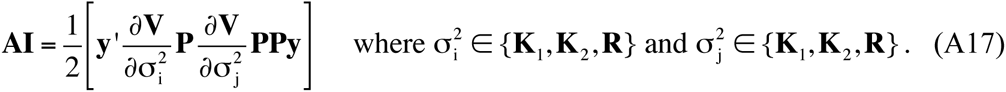

An implementation of the eigen-decomposition technique (Thompson and Shaw, 1990) in the log likelihood, the first derivatives (A16) and the **AI** matrix (A17) for the random regression is analogues to that in the multivariate linear mixed model, i.e. **ZAZ**’ in the matrix **V** can be replaced by **D**, thereby substantially reducing the computational complexity as shown in the previous section. As with the multivariate linear mixed model, the eigen-decomposition technique cannot be used for multiple genomic relationship matrices unless the additional relationship matrix is an identity matrix, which is to fit permanent environmental effects across different environments as in the model above.

In the average information algorithm for the random regression model based on the mixed model equations (Gilmour, et al., 2006; Meyer, 2007; Meyer and Hill, 1997), the log likelihood is written as

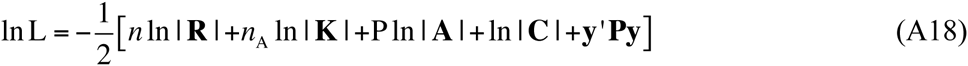

where *n* is the number of records, *n*_A_ is the number of individuals and **C** is the coefficient matrix in the mixed model equation (MME) (Henderson, 1975). It is noted that the classical AI algorithm based on equation (A18) is substantially different from what we propose here with different working variables.

#### 1.4. Convergence

With illegal or bad starting values, the REML process can have numerical problems. ASReml (Gilmour, et al., 2006), WOMBAT (Meyer, 2007) and GEMMA (Zhou and Stephens, 2014) use EM type algorithms in initial iterations to obtain a better set of starting values. This is mostly because if starting values are very different from the REML estimate, Newton type algorithms effectively update parameters that substantially depart from the current value, which occasionally results in updated values outside the legal parameter space. However, it is possible to prevent updated values from being in illegal states by moderating the magnitude of updates in the Newton type algorithm. The pseudo code of this moderating update is as follows:

~~~
Initial values assigned to parameter (**Θ**_0_);
*i=0*
Loop until REML estimate is found {
              *i* = *i* + 1
              **Δ_0_** is obtained from the first derivatives and **AI** matrix;
              **Θ_i_ = Θ_i–1_ + Δ_0_** (i.e. A8)
              *k* = 1;
              Log likelihood evaluation given the updated parameters **Θ_i_**
              If updated values are illegal
                            Update values are reduced by **Δ_k_** = **Δ_k–1_** × *f^k^*
                            **Θ_i_ = Θ_i–1_ + Δ*_k_***
                            *k* = *k* + 1
                            Goto log likelihood evaluation
              End if
}
~~~

We used an arbitrary value of *f* = 0.7 and observed no convergence and numerical problems with poor starting values (Table S8 and S9). Computational complexity for the moderating update requires only evaluation of the log likelihood, i.e. ~ *O*(*nt*^3^) in each update iteration (noted as *r*_1_ in Table S7, S8 and S9). The number of iterations needed to find a converged REML estimate is typically less than ten when using a moderating update strategy (Table S8 and S9). The moderating update strategy is also implemented in ASReml, but only in the first iteration round.

#### 1.5. Comparison of computational efficiency

We quantified the computational efficiency by comparing approximated computational complexity (Table S7). The computing time needed to find REML estimates was much lower for MTG2 and GEMMA than for ASReml and WOMBAT. MTG2 can be used for a wider range of statistical models than GEMMA, including multivariate linear mixed models, random regression models and multiple random effects models. GEMMA can only be used for a single random effect model in multivariate linear mixed models (Table S7). For random regression models or/and multiple random effects models, the computational efficiency for MTG2 (even without the eigen-decomposition) is considerably higher than that of ASReml and WOMBAT (Table S5 and S7). The moderating updating strategy further increases the relative computational efficiency of MTG2, resulting in a better performance of MTG2 compared with other methods. The advantage of MTG2 was larger when the number of traits increased (Table 1 and Table S4). We have checked the convergence behaviour for MTG2 with poor or good starting values, compared to that for ASReml, standard REML software using ten replicates of simulated phenotype data (Table S8 and S9). It is shown that the maximum log likelihood was perfectly agreed between MTG2 and ASReml and the process was converged well within a few iterations for both software whether using a multivariate (Table S8) or a random regression linear mixed model (Table S9). With good starting values, the number of iterations required for convergence reduced considerably for MTG2.

## 2. Supplementary Figures

**Figure S1.**
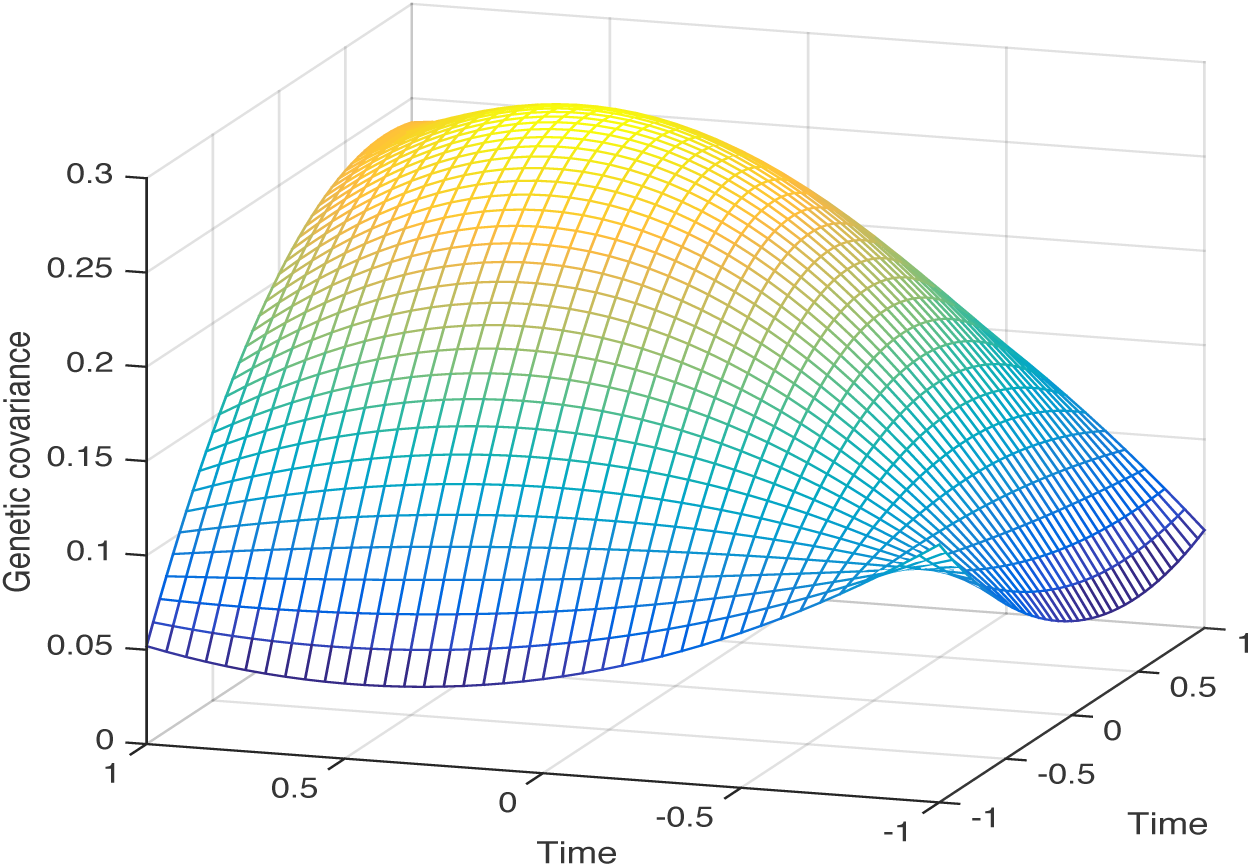
Genetic covariance pattern of glucose levels across time series after glucose injection from the random regression (see Supplementary Note). The order of Legendre polynomial is 3 for both genetic and residual variance components (see Table S3).

**Figure S2.**
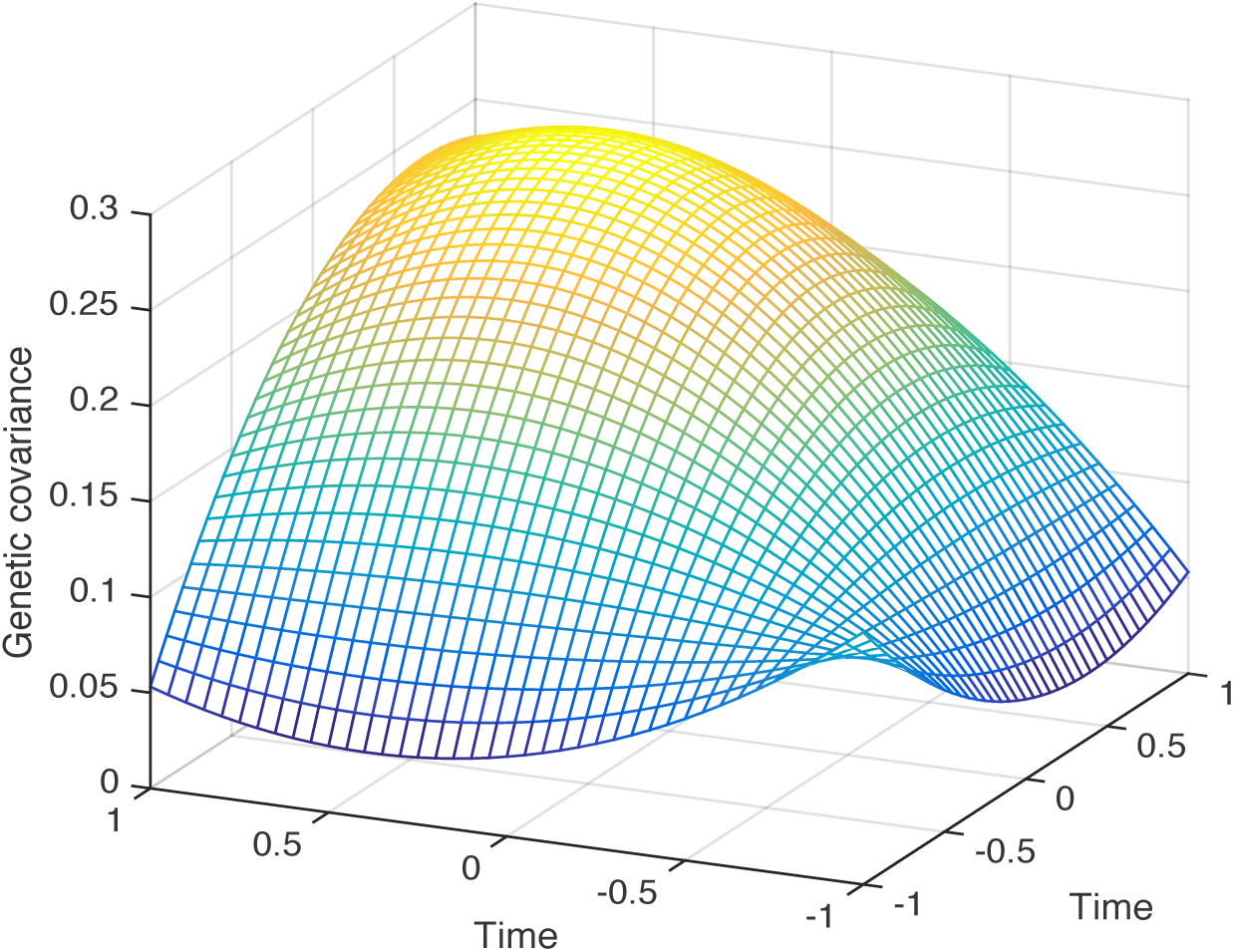
The same analysis as in Figure S1 except that the analysis was done without imputing missing phenotypes. This is to show the results from data with and without imputing missing phenotypic values are not much different.

## 3. Supplementary Tables

**Table S1.** Estimated SNP-heritability (h^2^) and genetic correlation (r_g_) between glucose level for each time after glucose injection and BMI when using the heterogeneous stock mice.

| Parameter | Traits | Estimation | SE |
| --- | --- | --- | --- |
| $h^2$ | Glucose 0 | 0.166 | 0.028 |
| $h^2$ | Glucose 15 | 0.147 | 0.026 |
| $h^2$ | Glucose 30 | 0.210 | 0.031 |
| $h^2$ | Glucose 75 | 0.239 | 0.032 |
| $h^2$ | BMI | 0.156 | 0.028 |
| $r_g$ | Glucose 0 and glucose 15 | 0.361 | 0.119 |
| $r_g$ | Glucose 0 and glucose 30 | 0.280 | 0.118 |
| $r_g$ | Glucose 15 and glucose 30 | 0.953 | 0.025 |
| $r_g$ | Glucose 0 and glucose 75 | 0.226 | 0.117 |
| $r_g$ | Glucose 15 and glucose 75 | 0.728 | 0.075 |
| $r_g$ | Glucose 30 and glucose 75 | 0.804 | 0.052 |
| $r_g$ | Glucose 0 and BMI | -0.004 | 0.137 |
| $r_g$ | Glucose 15 and BMI | -0.332 | 0.131 |
| $r_g$ | Glucose 30 and BMI | -0.153 | 0.128 |
| $r_g$ | Glucose 75 and BMI | 0.041 | 0.124 |

Estimates were essentially the same for the different REML estimation methods. The estimated SNP-heritability was moderate for the glucose level traits and BMI, ranging from 0.15 to 0.24. Among the glucose level straits, the genetic correlation between glucose 15 and 30 was the highest (0.95) while that between glucose 0 and 75 was the lowest (0.23). The genetic correlation between glucose levels within 30 minutes from the injection and BMI was negative, which was unexpected. However, the genetic correlation between glucose 75 and BMI was slight positive. More measurements after 75 minutes might be needed to find out the pattern. Further investigation is beyond the scope of this paper and warranted in a separate study.

**Table S2.**
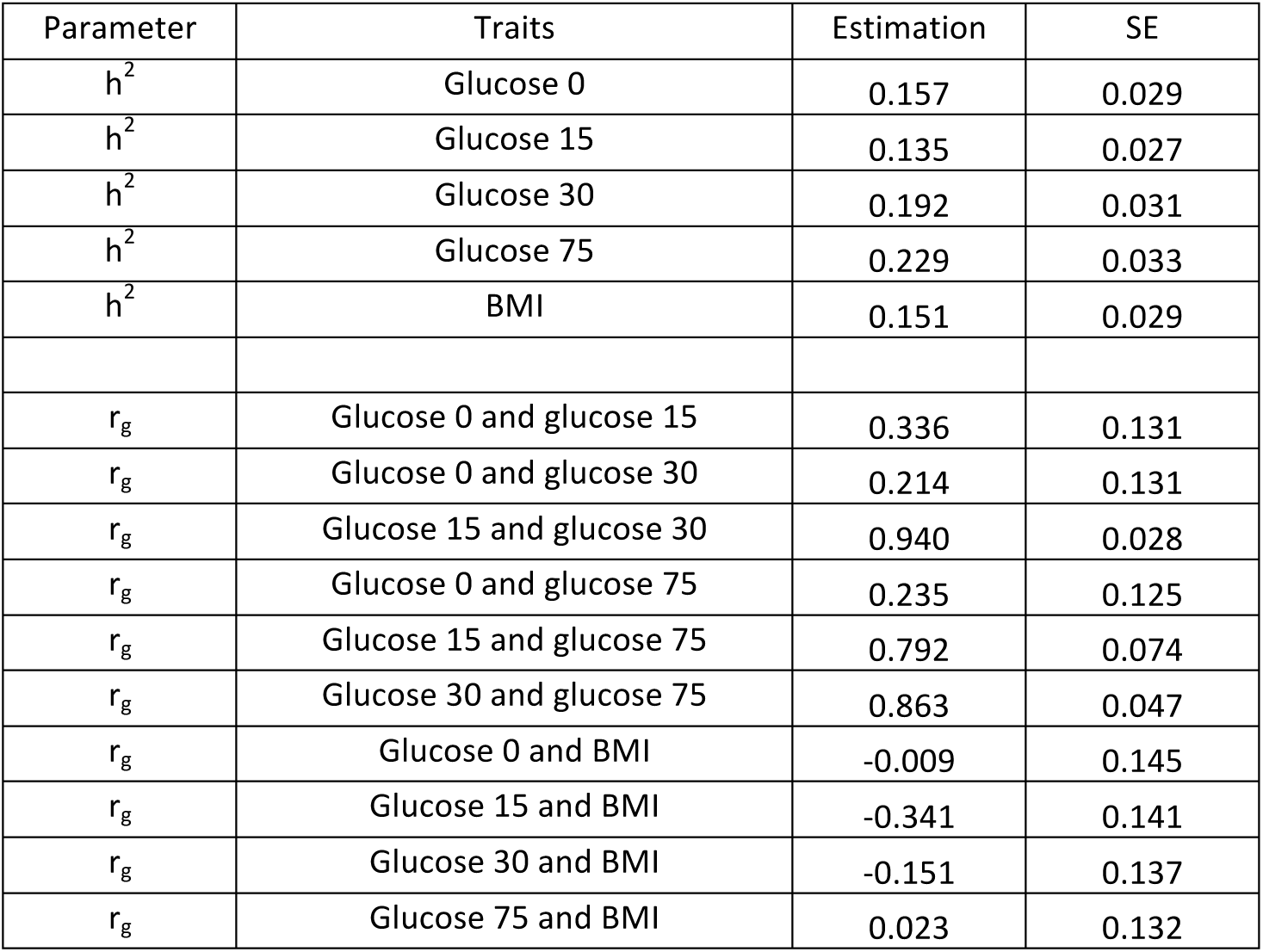
The same analysis as in Table S1 except that there was no phenotype imputation for missing values.

| Parameter | Traits | Estimation | SE |
| --- | --- | --- | --- |
| $h^2$ | Glucose 0 | 0.157 | 0.029 |
| $h^2$ | Glucose 15 | 0.135 | 0.027 |
| $h^2$ | Glucose 30 | 0.192 | 0.031 |
| $h^2$ | Glucose 75 | 0.229 | 0.033 |
| $h^2$ | BMI | 0.151 | 0.029 |
| $r_g$ | Glucose 0 and glucose 15 | 0.336 | 0.131 |
| $r_g$ | Glucose 0 and glucose 30 | 0.214 | 0.131 |
| $r_g$ | Glucose 15 and glucose 30 | 0.940 | 0.028 |
| $r_g$ | Glucose 0 and glucose 75 | 0.235 | 0.125 |
| $r_g$ | Glucose 15 and glucose 75 | 0.792 | 0.074 |
| $r_g$ | Glucose 30 and glucose 75 | 0.863 | 0.047 |
| $r_g$ | Glucose 0 and BMI | -0.009 | 0.145 |
| $r_g$ | Glucose 15 and BMI | -0.341 | 0.141 |
| $r_g$ | Glucose 30 and BMI | -0.151 | 0.137 |
| $r_g$ | Glucose 75 and BMI | 0.023 | 0.132 |

The estimates from the phenotypic data without imputing missing values (<10% for each trait) were very similar to those from the phenotypic data with imputing missing values (Table S1).

**Table S3.** Bayesian information criteria (BIC) for each model varying the number of order of Legendre polynomial in the random regression model.

|  |  | # Order for residual variance |  |  |
| --- | --- | --- | --- | --- |
|  |  | 1 | 2 | 3 |
| # Order for genetic variance | 1 | -7363.56 | -7708.08 | -8471.70 |
|  | 2 | -7700.28 | -7854.00 | -8606.18 |
|  | 3 | -7929.46 | -8110.98 | -8636.36 |
|  | 4 | -7939.51 | -8123.02 | -8623.00 |

It shows that the best model was with the order of 3 and 3 for genetic and residual variance component. We used BIC that is robust to type I errors although one can use AIC, likelihood ratio test or other variants of information criteria under their own justification. Figure S1 shows genetic covariance pattern of glucose levels across different time after intraperitoneal glucose injection, which was from the best model according to BIC.

**Table S4.**
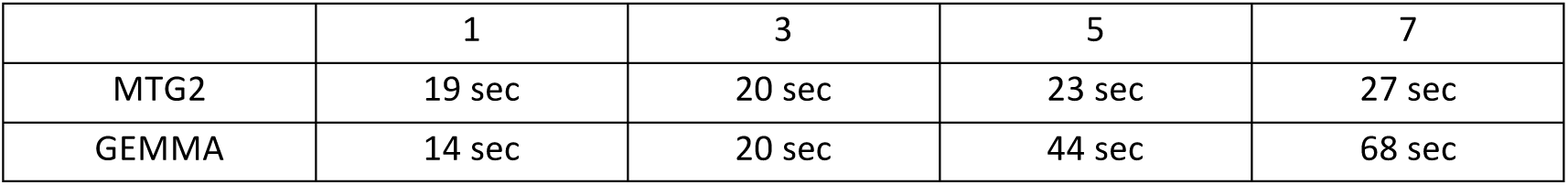
Computing time for MTG2 and GEMMA when using the ARIC cohort human data (N=7,263).

ASReml and WOMBAT were not used in this comparison. GEMMA was sightly faster than MTG2 when using a single trait model. With increasing number of traits, the computational efficiency for MTG2 was more increased than that for GEMMA. This would be expected given their computational complexity (Table S7). MTG2 and GEMMA took ~10 minutes for the eigen-decomposition, which only has to be done once per dataset then can then be reused for multiple analyses.

**Table S5.** Computing time for MTG2, ASReml and WOMBAT based on a single or multiple random effect models (i.e. multiple genetic covariance matrices) when using the heterogeneous stock mice data (N=1908).

|  | # Random effects |  |  |
| --- | --- | --- | --- |
|  | 1 | 2 | 3 |
|  | Multivariate linear mixed model (3 traits model) |  |  |
| MTG2 | 6min | 8min | 9min |
| ASReml | 210min | 1300 min | 4200min |
| WOMBAT | 9min | 62min | 130min |
|  | Random regression model<br>(# Order=2 for genetic and residual variance component) |  |  |
| MTG2 | 27min | 39min | 63min |
| ASReml | 82min | 470min | > 1000min |
| WOMBAT | 30min | 80min | 250min |

GEMMA was not included in these analyses because it cannot fit multiple random effects. MTG2 turns off the eigen-decomposition function and its performance was slowed down from the computing time of 1 second to 6 minutes for the three-trait linear mixed model, and from 2 seconds to 27 minutes for the random regression model with the order of 2 for both genetic residual variance components. In any case, the computational efficiency for MTG2 was the highest among the methods (Table S5). With increasing number of random effects (i.e. genetic covariance matrices), the relative computational efficiency for MTG2 was increased, compared to ASReml and WOMBAT. This would be expected because MTG2 was based on the direct AI algorithm using the variance covariance matrix of phenotypic observation and the size of this matrix is not affected by increasing number of random effects (see A(2)). However, MME-based AI algorithms (e.g. ASReml and WOMBAT) will slow down because the dimension of the MME increases with increasing number of random effects.

**Table S6.** Estimated SNP-heritability (h^2^) and genetic correlation (r_g_) between obesity and blood pressure traits when using ARIC cohort human data.

| Parameter | Traits | Estimation | SE |
| --- | --- | --- | --- |
| $h^2$ | BMI | 0.180 | 0.045 |
| $h^2$ | Triceps skinfold (TS) | 0.163 | 0.045 |
| $h^2$ | Waist girth (WG) | 0.164 | 0.045 |
| $h^2$ | Hip girth (HG) | 0.170 | 0.045 |
| $h^2$ | Waist to hip ratio (WHR) | 0.139 | 0.045 |
| $h^2$ | Systolic blood pressure (SP) | 0.193 | 0.045 |
| $h^2$ | Diastolic blood pressure (DP) | 0.201 | 0.045 |
| $h^2$ | Hypertension (HP) | 0.251<sup>a</sup> | 0.082 |
| $r_g$ | BMI and TS | 0.660 | 0.104 |
| $r_g$ | BMI and WG | 0.884 | 0.041 |
| $r_g$ | TS and WG | 0.614 | 0.122 |
| $r_g$ | BMI and HG | 0.741 | 0.069 |
| $r_g$ | TS and HG | 0.693 | 0.107 |
| $r_g$ | WG and HG | 0.856 | 0.056 |
| $r_g$ | BMI and WHR | 0.694 | 0.129 |
| $r_g$ | TS and WHR | 0.359 | 0.185 |
| $r_g$ | WG and WHR | 0.771 | 0.085 |
| $r_g$ | HG and WHR | 0.333 | 0.185 |
| $r_g$ | BMI and SP | 0.240 | 0.161 |
| $r_g$ | TS and SP | 0.159 | 0.175 |
| $r_g$ | WG and SP | 0.309 | 0.165 |
| $r_g$ | HG and SP | 0.159 | 0.171 |
| $r_g$ | WHR and SP | 0.289 | 0.183 |
| $r_g$ | BMI and DP | 0.242 | 0.159 |
| $r_g$ | TS and DP | 0.443 | 0.172 |
| $r_g$ | WG and DP | 0.129 | 0.172 |
| $r_g$ | HG and DP | 0.186 | 0.167 |
| $r_g$ | WHR and DP | -0.022 | 0.195 |
| $r_g$ | SP and DP | 0.768 | 0.075 |
| $r_g$ | BMI and HP | 0.270 | 0.192 |
| $r_g$ | TS and HP | 0.201 | 0.208 |
| $r_g$ | WG and HP | 0.324 | 0.199 |
| $r_g$ | HG and HP | 0.119 | 0.205 |
| $r_g$ | WHR and HP | 0.420 | 0.218 |
| $r_g$ | SP and HP | 0.970 | 0.138 |
| $r_g$ | DP and HP | 0.858 | 0.149 |

<sup>a</sup>Hypertension is a binary trait and the reported SNP-heritability is on the liability scale using the transformation (Dempster and Lerner, 1950; Lee, et al., 2011). The obesity traits (BMI, TS, WG, HG and WHR) and blood pressure traits (SP, DP and HP) were moderately heritable, with the estimated SNP-heritability ranging from 0.14 to 0.20. Among the obesity traits, there were positive genetic correlations (0.33 – 0.88). For the blood pressure traits, there were also strong positive genetic correlations (0.77 – 0.97). Between the obesity and blood pressure traits, all of the estimated genetic correlations were positive except one between WHR and DP (-0.02). The genetic correlation between TS and DP was the highest (0.44), followed by that between WHR and HP (0.42).

**Table S7.** Approximated computational complexity of the methods

|  | Multivariate linear mixed model | Random regression model |
| --- | --- | --- |
|  | With a single genetic covariance matrix |  |
| MTG2 | $O(n^3 + n^2t + n^2c + r_1nt^3 + r_2nct^6)$ | $O(n^3 + n^2p + n^2c + r_1np^3 + r_2ncp^2k^4)$ |
| GEMMA | $O(n^3 + n^2t + n^2c + r_3nc^2t^2 + r_2nc^2t^6)$ | NA |
| ASReml | $O(r_3n^3(t+c)^3 + r_2n^3t^7)$ | $O(r_3n^3(p+c)^3 + r_2n^3p^3k^4)$ |
| WOMBAT | $O(r_3n^3(t+c)^3 + r_2n^3t^7)$ | $O(r_3n^3(p+c)^3 + r_2n^3p^3k^4)$ |
| | With multiple genetic covariance matrices ($m>1$) | |
| MTG2 | $O(r_1n^3t^3 + r_2n^3t^7m^4)$ | $O(r_1n^3p^3 + r_2n^3p^3k^4m^4)$ |
| GEMMA | NA | NA |
| ASReml | $O(r_3n^3m^3(t+c)^3 + r_2n^3t^7m^7)$ | $O(r_3n^3m^3(p+c)^3 + r_2n^3p^3k^4m^7)$ |
| WOMBAT | $O(r_3n^3m^3(t+c)^3 + r_2n^3t^7m^7)$ | $O(r_3n^3m^3(p+c)^3 + r_2n^3p^3k^4m^7)$ |

*n* is the number of individuals, *t* is the number of traits, *c* is the number of fixed effects, *r*_1_ is the number of iterations for the moderating update, *r*_2_ is the number of iteration for Newton type algorithm, *r*_3_ is the number of iterations for EM-type algorithm, *p* is the number of environments, and *m* is the number of genetic covariance matrices (i.e. GRM).

**Table S8.** Obtained log likelihood, the number of iterations and convergence when using MTG2 and ASReml with three-trait linear mixed model for the WTCCC heterogeneous mice stock.

| Replicates | MTG2 |  |  |  |  | ASReml<sup>c</sup> |  | Converged |
| --- | --- | --- | --- | --- | --- | --- | --- | --- |
|  | Likelihood | # iteration |  |  |  | Likelihood | # iteration<br>(r<sub>2</sub><sup>e</sup>) |  |
|  |  | Poor starting<br>value<sup>a</sup> |  | Good starting<br>value<sup>b</sup> |  |  |  |  |
|  |  | r<sub>1</sub><sup>d</sup> | r<sub>2</sub><sup>e</sup> | r<sub>1</sub><sup>d</sup> | r<sub>2</sub><sup>e</sup> |  |  |  |
| 1 | -4424.15 | 2 | 6 | 0 | 3 | -4424.15 | 9 | yes |
| 2 | -4378.5 | 11 | 10 | 0 | 4 | -4378.5 | 8 | yes |
| 3 | -4375.63 | 3 | 6 | 0 | 4 | -4375.63 | 8 | yes |
| 4 | -4326.18 | 3 | 6 | 0 | 3 | -4326.18 | 8 | yes |
| 5 | -4354.97 | 3 | 6 | 0 | 3 | -4354.97 | 15 | yes |
| 6 | -4411.81 | 2 | 6 | 0 | 4 | -4311.81 | 14 | yes |
| 7 | -4093.76 | 2 | 5 | 0 | 3 | -4093.76 | 9 | yes |
| 8 | -4216.42 | 2 | 6 | 0 | 4 | -4216.42 | 10 | yes |
| 9 | -4119.63 | 2 | 5 | 0 | 3 | -4119.63 | 8 | yes |
| 10 | -4218.67 | 2 | 6 | 0 | 4 | -4218.67 | 10 | yes |

Ten replicates of phenotypic data were used to assess the performance. Simulation was based on multivariate normal variables reflecting the variance and covariance structure estimated in the random regression model for the glucose traits (Figure S1). ^a^Arbitary values were assigned, e.g. genetic and residual variances were assigned as half of the total phenotypic variance and covariances was assigned as zero. ^b^True simulated parameters were used. ^c^ASReml hardly used EM algorithm unless there were numerical problems. ^d^*r*_1_ is the number of iterations for the moderating update (see Table S7). ^e^*r*_2_ is the number of iteration for Newton type algorithm (see Table S7).

**Table S9.** Obtained log likelihood, the number of iterations and convergence when using MTG2 and ASReml with random regression model with Legendre polynomial order of 3 for genetic and residual variance components. The heterogeneous mice stock was used with simulated phenotypic data.

| Replicates | MTG2 |  |  |  |  | ASReml |  | Converged |
| --- | --- | --- | --- | --- | --- | --- | --- | --- |
|  | Likelihood | # iteration |  |  |  | Likelihood | # iteration (r<sub>2</sub>) |  |
|  |  | Poor starting value |  | Good starting value |  |  |  |  |
|  |  | r<sub>1</sub> | r<sub>2</sub> | r<sub>1</sub> | r<sub>2</sub> |  |  |  |
| 1 | -8324.47 | 0 | 9 | 0 | 3 | -8324.47 | 7 | yes |
| 2 | -8311.27 | 0 | 8 | 0 | 3 | -8311.27 | 7 | yes |
| 3 | -8259.34 | 0 | 8 | 0 | 3 | -8259.34 | 7 | yes |
| 4 | -8325.44 | 0 | 8 | 0 | 3 | -8325.44 | 7 | yes |
| 5 | -8315.36 | 0 | 8 | 0 | 3 | -8315.36 | 7 | yes |
| 6 | -8292.39 | 0 | 8 | 0 | 3 | -8392.39 | 7 | yes |
| 7 | -8367.17 | 0 | 8 | 0 | 3 | -8367.17 | 7 | yes |
| 8 | -8363.37 | 0 | 8 | 0 | 3 | -8363.37 | 7 | yes |
| 9 | -8329.45 | 0 | 8 | 0 | 3 | -8429.45 | 8 | yes |
| 10 | -8371.66 | 0 | 8 | 0 | 3 | -8371.66 | 7 | yes |

## 4. Acknowledgments

The Atherosclerosis Risk in Communities Study is carried out as a collaborative study supported by National Heart, Lung, and Blood Institute contracts HHSN268201100005C, HHSN268201100006C, HHSN268201100007C, HSN268201100008C, HHSN268201100009C, HHSN268201100010C, HSN268201100011C, and HHSN268201100012C). The authors thank the staff and participants of the ARIC study for their important contributions. Funding for CARe genotyping was provided by NHLBI Contract N01-HC-65226. Funding for GENEVA was provided by National Human Genome Research Institute grant U01HG004402 (E. Boerwinkle).

